# Gaussian Accelerated Molecular Dynamics in GROMACS

**DOI:** 10.64898/2026.08.10.743837

**Authors:** Yuefeng Yang

**Author notes:** Corresponding authors: Yuefeng Yang.

## Abstract

Gaussian accelerated molecular dynamics (GaMD) enhances conformational sampling by adding a smooth boost potential without requiring predefined collective variables, but an engine-integrated implementation has not been available in GROMACS. Here, we implement total-, dihedral-, and dual-boost GaMD in GROMACS 2025.4, including staged energy-statistics collection, GPU-based bias evaluation and force scaling, restart support, and outputs required for cumulant-based free-energy reweighting. The implementation was evaluated using four benchmark systems spanning conformational free energies, protein folding, and ligand recognition. For alanine dipeptide, a reweighted 100 ns GaMD trajectory recovered the major free-energy basins and rotational barriers in overall agreement with a 1000 ns conventional MD simulation. For chignolin and TC5b, all three independent trajectories for each system sampled native-like folded states from extended conformations within 300 ns and 1 μs, respectively; the best TC5b structure had a minimum backbone RMSD of 0.03 nm from the experimental structure. In the benzene–T4 lysozyme system, two of five independent 500 ns trajectories captured both ligand binding and dissociation, yielding a bound pose with a minimum ligand RMSD of 0.06 nm from the crystal structure. Across all four systems, the boost-potential distributions were approximately Gaussian, and second-order cumulant reweighting resolved the expected conformational and binding free-energy basins. These results demonstrate that GROMACS-GaMD provides a practical, GPU-enabled, collective-variable-free enhanced-sampling framework for biomolecular free-energy calculations, protein folding, and ligand-binding studies.

## INTRODUCTION

Atomistic molecular dynamics (MD) simulations have become an indispensable tool for linking molecular structure to thermodynamic properties and dynamic mechanisms. Nevertheless, the fundamental limitation of conventional MD remains its restricted accessible time scale: many biologically relevant conformational transitions, folding events, and binding or unbinding processes are separated by free-energy barriers and therefore cannot be adequately sampled within routine simulation windows. Improving the sampling efficiency of such rare events while retaining atomistic resolution thus remains a central challenge in molecular simulation and has motivated the continued development of enhanced-sampling methods for complex molecular systems.^1,2^

Among the available enhanced-sampling strategies, Gaussian accelerated molecular dynamics (GaMD) is particularly attractive from a methodological standpoint.^1,2^ Unlike biasing approaches that rely on predefined collective variables, GaMD reduces effective barriers by directly modifying the potential-energy surface. Specifically, when the system potential falls below a prescribed threshold, a smooth boost potential is applied to facilitate transitions among metastable states.^1^ Meanwhile, the threshold and force constant are constrained by the statistical properties of the potential energy so that the boost potential remains approximately Gaussian, thereby enabling postprocessing reweighting through cumulant expansion.^1,2^ Owing to this design, GaMD can substantially enhance configurational sampling without requiring prior specification of reaction coordinates, while still permitting practical recovery of free-energy information. This feature is especially valuable for biomolecular systems with complex degrees of freedom and transition pathways that are difficult to define a priori.^1,2^

Although GaMD is now a mature methodology, its software support across major MD engines remains uneven. AMBER currently provides the most comprehensive publicly available GaMD family, including the standard formulation as well as multiple selective variants.^3–5^ NAMD also offers official implementations of total-, dihedral-, and dual-boost modes, whereas related efforts in OpenMM have demonstrated that GaMD can be ported to modern modular and extensible simulation frameworks.^6,7^ By contrast, PLUMED is primarily organized around collective-variable-based biasing methods, and current public documentation does not identify a dedicated official GaMD action.^8^ Likewise, although GROMACS offers a rich enhanced-sampling ecosystem, including AWH, replica exchange, pull-code methods, Colvars, and a limited PLUMED interface, current official documentation does not provide a native GaMD module (Table 1).^9^ In other words, while the scientific value of GaMD has been well established, its availability in GROMACS, one of the most widely used high-performance MD engines, has remained limited.

**Table 1.**
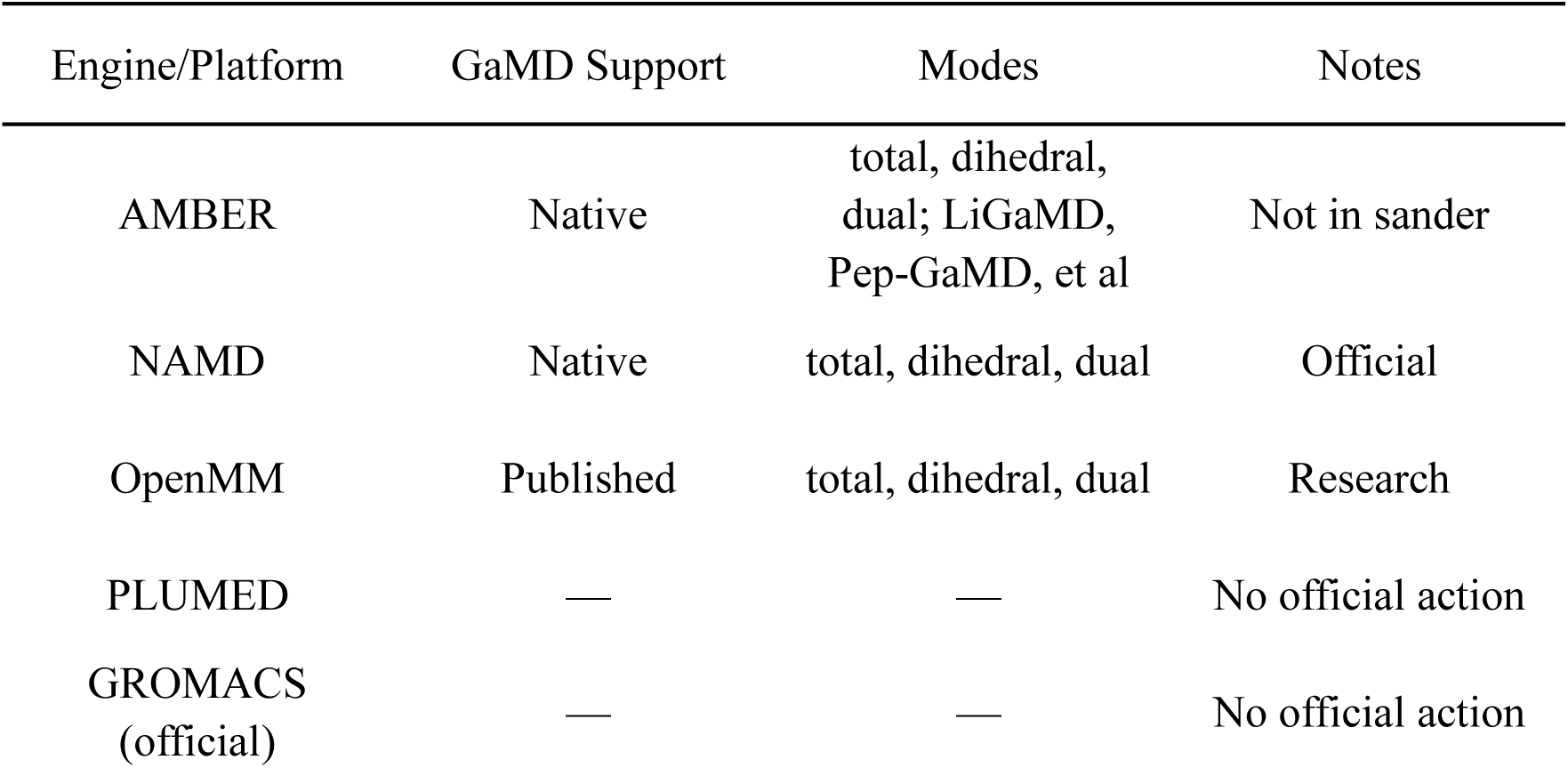
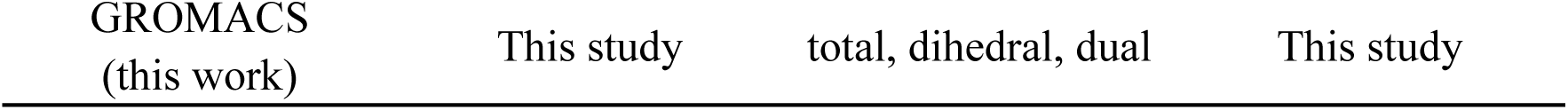
Publicly documented GaMD support across major molecular dynamics engines.

This gap makes the implementation of GaMD in GROMACS important from both scientific and software-engineering perspectives. From the standpoint of applications, GROMACS combines broad community adoption with mature parallel performance and a robust simulation ecosystem, making it a natural platform for extending access to collective-variable-free enhanced sampling. From the standpoint of implementation, however, this effort is not merely a straightforward feature port. It requires engine-level treatment of several key issues, including reliable access to the relevant energy terms, strict consistency between bias forces and reported energies, continuous updating of running potential-energy statistics, and reproducible algorithmic behavior across restart conditions, parallel settings, and heterogeneous hardware environments. Only when these theoretical and engineering requirements are simultaneously satisfied can GaMD function within GROMACS as a verifiable, reusable, and scientifically meaningful enhanced-sampling method.

Against this background, we implemented GaMD in GROMACS and established a practical workflow encompassing bias construction, force scaling, GPU implementation, and subsequent validation and benchmarking. This implementation not only fills a practical gap in the enhanced-sampling capabilities of GROMACS by providing native GaMD support, but also lays the groundwork for future extensions to selective or hybrid GaMD variants. The remainder of this paper is organized as follows. We first present the theoretical basis of GaMD and the design of its implementation in GROMACS, then describe validation and benchmark results, and finally discuss the limitations of the present implementation and directions for future development.

## METHODS

### Theory

Gaussian accelerated molecular dynamics (GaMD) is an unconstrained enhanced-sampling method that accelerates barrier crossing by adding a smooth harmonic boost potential to the underlying potential-energy surface. Unlike biasing schemes that require predefined collective variables, GaMD acts directly on the potential energy and is therefore well suited to systems for which the relevant transition coordinates are not known a priori. A defining feature of the method is that the boost potential is constructed to remain approximately Gaussian, which in turn allows unbiased free-energy profiles to be recovered by cumulant-based reweighting.^2^

For consistency with the present implementation, the potential energy subject to boosting is denoted by *V* (**r**), where **r** represents the full set of atomic coordinates. The GaMD-modified potential is written as

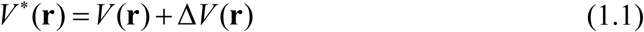

where the boost potential Δ*V* (**r**) is applied only when the instantaneous potential energy falls below a prescribed threshold energy *E*:

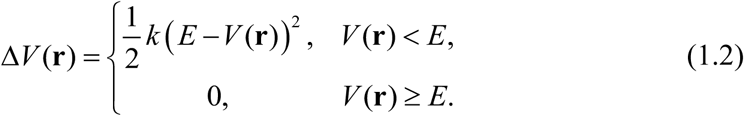

Here, *k* is the harmonic force constant controlling the magnitude of the boost. This construction selectively raises low-energy regions of the potential-energy surface, thereby reducing effective energy barriers and facilitating transitions among metastable states. Depending on the chosen boost mode, *V* (**r**) may represent the total potential energy, a dihedral-energy term, or another designated energy component. For multi-boost formulations, the total GaMD bias can be expressed as the sum of individual boost contributions,

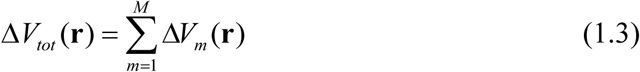

so that the biased potential becomes

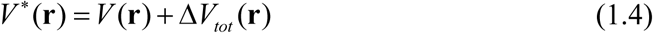

To ensure that the boosted potential remains physically meaningful and amenable to reweighting, the threshold energy *E* and force constant *k* are determined from the statistical properties of the potential energy collected during the preparatory stage. Let *V*_min_, *V*_max_, *V*_avg_, and *σ_V_* denote the minimum, maximum, average, and standard deviation of the potential energy, respectively. GaMD parameterization is constrained such that the ordering of potential energies is preserved after boosting (*V_1_*\* < *V_2_*\*), the boosted potential energy differences remain smaller than the corresponding original potential energy differences (*V_2_*\* −*V_1_* * < *V_2_*−*V_1_*). These requirements lead to the condition

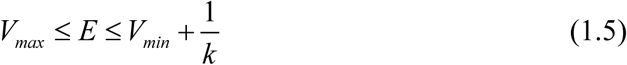

According to the inequality constraints, the force constant can be expressed as

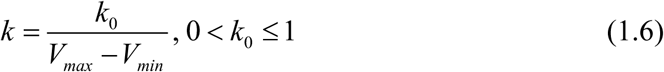

In addition, the standard deviation of the boost potential is required to remain below a user-defined upper bound *σ* _0_:

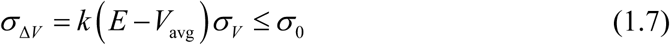

In practice, two commonly used parameter-selection schemes are employed. In the first scheme, the threshold energy is taken at its lower bound,

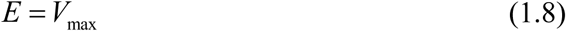

which yields

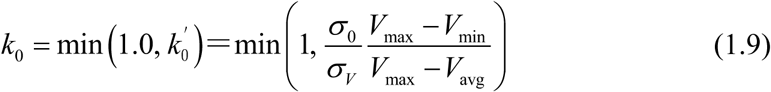

In the second scheme, the threshold energy is placed at its upper bound,

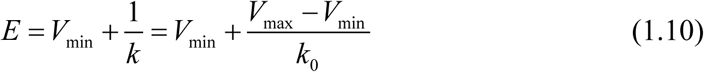

Under this choice, one obtains

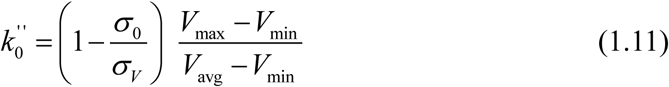

If 0 < *k_0_* ^’’^ ≤ 1, then 0 < *k_0_* ^’’^ is adopted; otherwise, the parameterization reverts to the *E* = *V*_max_ scheme. These expressions formalize the balance between sampling acceleration and reweighting fidelity: a larger boost may enhance barrier crossing, but excessive broadening of the boost distribution compromises the accuracy of free-energy reconstruction.

The GaMD bias enters the equations of motion through the biased force. Differentiating the modified potential gives

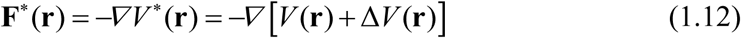

For configurations satisfying *V* (**r**) < *E*, this becomes

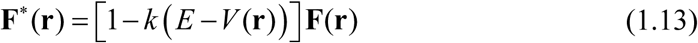

where

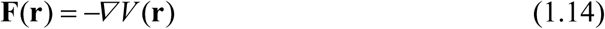

Thus, GaMD can be interpreted as an energy-dependent scaling of the original force. This form is particularly convenient for practical implementation because the bias can be applied through a local force-scaling operation once the instantaneous energy and the corresponding boost factor have been determined.

Because GaMD generates a biased ensemble, postprocessing reweighting is required to recover unbiased thermodynamic observables. Let *A* denote a reaction coordinate or any collective observable of interest. The unbiased probability distribution *p* _(_*A_j_* _)_ for bin *A_j_* is related to the biased distribution *p\**_(_*A_j_*_)_ through

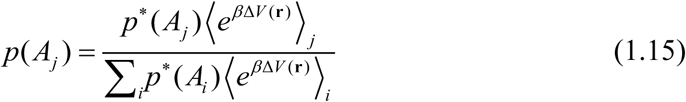

where

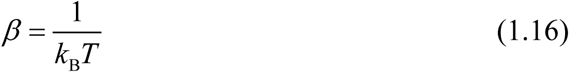

The corresponding unbiased free energy is then given by

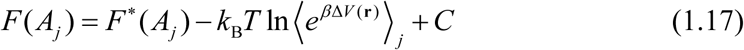

with *C* being an arbitrary constant that sets the zero of free energy.

A central assumption of GaMD is that the boost potential is approximately Gaussian distributed. Under this assumption, the exponential reweighting factor can be approximated by a cumulant expansion,^10,11^

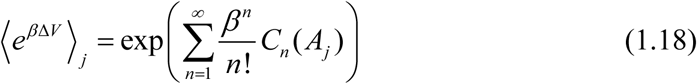

where *C_n_* _(_*A_j_* _)_ is the *n*-th cumulant of the boost-potential distribution within bin *A_j_*. Retaining terms up to second order yields

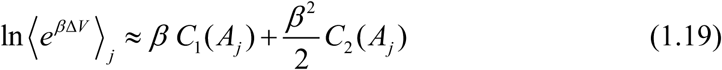

with

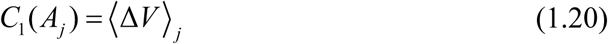

and

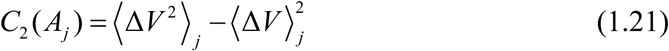

Accordingly, the unbiased free energy can be approximated as

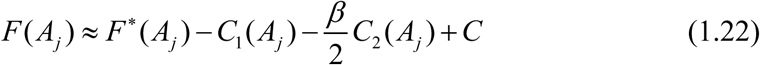

This second-order approximation provides the theoretical basis for practical GaMD reweighting and is reliable when the boost distribution remains narrow and close to Gaussian, which is precisely why the parameter *σ* _0_ plays a central role in the method.

In a typical GaMD workflow, a short conventional MD stage is first used to collect potential-energy statistics and estimate *V*_min_, *V*_max_, *V*_avg_, and *σ_V_*. A subsequent bias-equilibration stage applies the initial boost while continuing to update these quantities, after which production GaMD simulations are performed and the accumulated boost potential is recorded for reweighting analysis. In this way, GaMD combines barrier reduction during sampling with a statistically controlled route for approximating the corresponding unbiased free-energy landscape.

### Implementation

GaMD was implemented as an engine-level extension to GROMACS 2025.4^9^ rather than as an external wrapper or postprocessing utility. The method was enabled through the gamd keyword in the mdp file, whereas GaMD-specific control parameters were provided separately in gamd.in, including the boost mode (igamd), threshold-energy options (iE, iEP, and iED), stage lengths (ntcmdprep, ntcmd, ntebprep, and nteb), the averaging window (ntave), and the output-control parameters reweight_nst, para_nst, sigma0P, and sigma0D. The input values of sigma0P and sigma0D were specified in kcal/mol and converted internally to kJ/mol.

The implementation followed the standard staged GaMD protocol,^2^ comprising cMD preparation, cMD statistics collection, boost equilibration preparation, boost equilibration statistics, and production. During the cMD statistics stage, extrema, mean values, and standard deviations of the target energies were accumulated and used to initialize the threshold energy *E*, the normalized force constant *k*_0_, and the effective force constant *k*. During boost equilibration, the bias potential was applied while the running statistics and acceleration parameters continued to be updated. Production GaMD simulations were then carried out using the final parameter set.

At the implementation level, the total-boost and dihedral-boost components were handled separately. Nonbonded, PME, bonded, and update calculations were offloaded to the GPU through the native GROMACS framework, while dedicated GaMD GPU kernels evaluated the boost factors and applied the corresponding force corrections. During fixed-parameter production, energies and forces remained on the GPU whenever possible, reducing host–device transfers and enabling CUDA Graph acceleration. Stage control, the collection of adaptive statistics, parameter updates, restart management, and the generation of reweighting output were handled on the host. Production simulations were performed using gmx mdrun-deffnm gamd-ntmpi 1 - ntomp X-pin on-nb gpu-pme gpu-bonded gpu-update gpu.

To support restart and postprocessing analysis, dedicated GaMD state and output files were introduced to preserve statistical state, record parameter evolution, and write the energy and weight terms required for reweighting. These outputs included the evolution of *V*_max_, *V*_min_, *V*_avg_, *σ_V_*, *E*, *k*_0_, and *k*, together with the unboosted energies, boost energies, and associated force-weight terms. All GaMD results reported here were obtained using the modified GROMACS 2025.4 build on an Intel i9-14900KF CPU and an NVIDIA GeForce RTX 4080 GPU.

### Simulation Protocols and Benchmarks

To evaluate the applicability and reliability of the GaMD implementation developed in this work in GROMACS, we selected four representative benchmark systems: alanine dipeptide in explicit solvent, the fast-folding peptide chignolin^12^and TC5b,^13^ and benzene binding to T4 lysozyme.^14^ These four systems probe three distinct aspects of enhanced-sampling performance, namely the accuracy of free-energy reweighting in small molecule/peptide systems, the ability to enhance conformational sampling during protein folding, and the characterization of ligand-binding pathways and bound states in protein-ligand recognition.

For all four systems, simulation systems for both conventional MD and GaMD were constructed using CHARMM-GUI.^15^ After steepest-descent energy minimization, each system was equilibrated in the NVT ensemble, and all subsequent cMD and GaMD simulations were carried out in the NPT ensemble. The CHARMM36m force field^16^ was used for the biomolecular components, and TIP3P was used as the water model.^17^ Benzene was parameterized with GAFF2,^18^ with partial charges assigned using the AM1-BCC method.^19^ Long-range electrostatic interactions were treated with Particle Mesh Ewald (PME).^20^ A van der Waals cutoff of 1.2 nm was used, and the neighbor list was updated every 20 steps. Bonds involving hydrogen atoms were constrained using LINCS,^21^ allowing a 2 fs integration time step. Temperature was maintained at 300 K using the v-rescale^22^ thermostat with a coupling time constant of 1 ps, and pressure was maintained at 1 bar using the c-rescale^23^ barostat with a coupling time constant of 5 ps. For all GaMD simulations, potential energies were evaluated at every MD step by setting nstcalcenergy = 1. All systems were simulated in the dual-boost mode, in which boost potentials were applied to both the total potential energy and the dihedral energy. In addition, the threshold energies were set to the lower bound, and both sigma0P and sigma0D were set to the default value of 6 kcal/mol. With these sigma0 settings, the resulting k_0_ values were both 1 after equilibration with the GaMD boost potential.

The alanine dipeptide system was used to assess the accuracy of GaMD free-energy reweighting in a small system. Because the conformational landscape of alanine dipeptide is primarily described by the backbone dihedral angles *ϕ* and *ψ*, this benchmark focused on whether GaMD could recover the major low-energy basins and the rotational barriers on the two-dimensional free-energy surface. The system was solvated in a 3.0 nm × 3.0 nm × 3.0 nm box, giving a total of 2252 atoms. In the GaMD simulations, the cMD stage was run for 3 ns, followed by 6 ns of boost equilibration and 100 ns of GaMD production. For comparison, a 1000 ns conventional MD simulation was carried out. In the subsequent analysis, the *ϕ* and *ψ* dihedral-angle distributions were extracted, and second-order cumulant expansion was used to reweight the GaMD trajectories and construct the two-dimensional (*ϕ*,*ψ*) potential of mean force (PMF).

The chignolin system was used to evaluate the ability of the present implementation to enhance conformational sampling during protein folding. The unfolded initial conformation was generated using AmberTools,^24^ while the folded reference structure was taken from PDB entry 1UAO.^12^ The system was solvated in a 5.3 nm × 5.3 nm × 5.3 nm box, with a total of 14,599 atoms. In the GaMD simulations, the cMD stage was run for 5 ns, followed by 30 ns of boost equilibration and 300 ns of GaMD production for each trajectory. Three independent GaMD production runs were performed using different initial velocity seeds. In the subsequent analysis, the *C_α_* RMSD relative to the folded reference structure and the radius of gyration *R_g_* were used as the primary reaction coordinates, and the corresponding two-dimensional PMF was constructed to identify the folded, intermediate, and unfolded conformational basins. To further demonstrate the practical applicability of the GaMD algorithm, we evaluated it using the more complex TC5b system. The initial conformation was also generated using AmberTools, and the system contained 68,235 atoms. The experimental structure with PDB ID 1L2Y was used as the folded reference structure.^13^ For each GaMD trajectory, a 10 ns cMD stage was followed by 50 ns of GaMD equilibration and a 1,000 ns GaMD production stage. Three independent GaMD simulations were performed using different random seeds for the initial velocities. In the subsequent analysis, a two-dimensional PMF profile was constructed using the same reaction coordinates as those employed for the chignolin system. These two benchmarks were designed to determine whether GaMD could capture folding and unfolding events on the hundred-nanosecond timescale, enhance configurational sampling relative to conventional MD, and resolve distinct free-energy basins corresponding to folded, intermediate, and unfolded states.

The benzene-T4 lysozyme system was used to evaluate the ability of the present implementation to enhance sampling of protein-ligand binding processes. The initial structure was based on the benzene-bound T4 lysozyme complex, using PDB entry 181L^14^ as the structural reference. In the starting configuration for production simulations, the benzene molecule occupying the binding pocket was removed, and 10 benzene molecules were randomly placed in the solvent. The system was solvated in a 7.6 nm × 7.6 nm × 7.6 nm box, with a total of 44,320 atoms. In the GaMD simulations, the cMD stage was run for 5 ns, followed by 30 ns of boost equilibration. Five independent GaMD production trajectories were then carried out, each with a length of 500 ns. Among these, two trajectories sampled benzene binding and dissociation events. In the subsequent analysis, ligand RMSD and the protein-ligand contact number *N*_contact_ were used to characterize the binding process, and a two-dimensional PMF was constructed based on these coordinates to distinguish the unbound, intermediate, and bound states. This benchmark was designed to assess whether GaMD could capture the pathways of benzene entering and leaving the binding pocket on an accessible timescale, recover binding poses consistent with the crystal structure, and resolve key intermediate states along the binding process.

## RESULTS

### Alanine Dipeptide

In the alanine dipeptide example, the PMF is calculated using *ϕ* and *ψ* as the two collective variables (CVs, Figure 1A). The probability distribution of the boost potential, Δ*V*, is unimodal and approximately Gaussian, with its maximum centered at approximately 11–12 kcal/mol, and an anharmonicity of 8.91×10^-3^ (Figure 1B). This value is consistent with the low anharmonicity previously reported for alanine dipeptide in the original NAMD-GaMD implementation, indicating that the boost-potential distribution in the present simulations remains sufficiently close to Gaussian to justify reweighting based on the second-order cumulant expansion.^6^ As shown in Figure 1C– D, the 100 ns GaMD production simulation, after reweighting using the second-order cumulant expansion, reproduced both the one-dimensional and two-dimensional free-energy profiles of alanine dipeptide with good overall agreement relative to the 1000 ns cMD reference.

**Figure 1.**
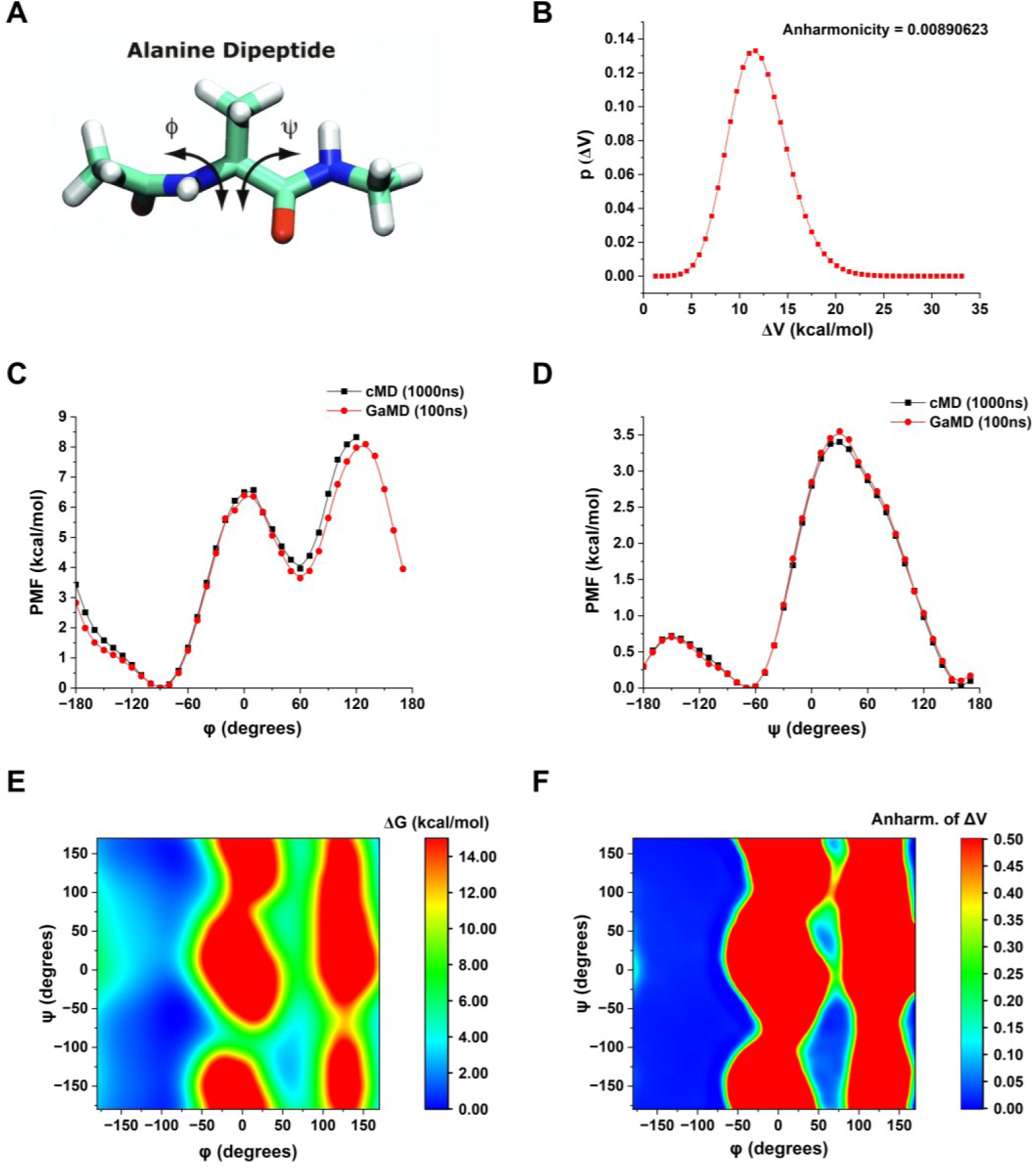
Validation results of GaMD for the alanine dipeptide system. (A) Molecular structure of alanine dipeptide, with the backbone dihedral angles *ϕ* and *ψ* indicated. (B) Probability distribution of the boost potential Δ*V* during the 100 ns GaMD production simulation. The corresponding anharmonicity is 8.9×10^-3^, indicating that the boost-potential distribution remains close to Gaussian. (C, D) One-dimensional PMFs projected onto the *ϕ* and *ψ* dihedral angles, respectively, comparing the reweighted 100 ns GaMD results with the 1000 ns cMD reference. (E) Two-dimensional reweighted PMF constructed from the *ϕ* and *ψ* dihedral angles. (F) Local anharmonicity map of the boost potential in the (*ϕ*,*ψ*) plane.

The one-dimensional PMF projected onto the *ϕ* dihedral angle (Figure 1C) shows that the reweighted GaMD result closely overlaps with the 1000 ns cMD profile and successfully reproduces the principal barrier near *ϕ* ≍ 0, with a height of approximately 6.3–6.5 kcal/mol. This value agrees well with the forward *ϕ*-rotation barrier of 6.07 kcal/mol reported by Miao et al. for alanine dipeptide in explicit solvent.^25^ From the one-dimensional projection, the dominant low-free-energy basin along *ϕ* spans approximately −120 to −80, with a minimum near *ϕ* ≍ −100. In addition, a secondary local minimum is observed at *ϕ* ≍ 50°–70°. Beyond the main barrier, the positive-*ϕ* region exhibits a higher barrier, peaking at approximately *ϕ* ≍ 110°–130° with a height of about 8.0 kcal/mol, indicating that conformations in the positive-*ϕ* region are substantially less thermodynamically accessible than those in the dominant negative-*ϕ* basin under the present force-field conditions. Compared with GaMD, cMD did not sample the corresponding conformations once *ϕ* exceeded approximately 120°.

The one-dimensional PMF projected onto the *ψ* dihedral angle (Figure 1D) likewise shows good agreement between cMD and reweighted GaMD. In the present system, the dominant low-free-energy basin is located at approximately *ψ* ≍ −70° to −50°, with the minimum near *ψ* ≍ −60°. A second pronounced low-energy basin is also observed in the region *ψ* ≍ 140°–170°. The principal barrier occurs near *ψ* ≍ 20°–50°, with a height of approximately 3.5 kcal/mol.

The two-dimensional (*ϕ*,*ψ*) PMF (Figure 1E) further shows that the major low-free-energy regions are concentrated on the negative-*ϕ* side, forming a relatively broad low-energy channel, whereas the positive-*ϕ* region remains globally less favorable. It should be emphasized that the present two-dimensional free-energy surface does not correspond exactly to the alanine dipeptide free-energy landscapes reported in the early AMBER-GaMD and NAMD-GaMD studies.^2,6^ Such differences are expected, because those early implementations were based on systems parameterized with an AMBERff99,^26^ whereas the present simulations were performed using CHARMM36m. In this respect, the free-energy landscape obtained here is more consistent with the more recent NAMD-GaMD tutorial than with the original AMBER-and NAMD-GaMD benchmark results.^27^

Finally, the local anharmonicity map (Figure 1F) shows that the anharmonicity of Δ*V* remains low within the principal low-free-energy basins, whereas elevated anharmonicity is mainly observed in the positive-*ϕ* region and other high-free-energy, sparsely sampled areas. This indicates that the reweighted free-energy estimates are most reliable in the thermodynamically dominant conformational regions, while greater caution is warranted in high-energy regions with limited sampling. Taken together, these results demonstrate that the GaMD implementation developed here in GROMACS is capable of recovering the major free-energy features of alanine dipeptide using a sampling time an order of magnitude shorter than that required for cMD. At the same time, because the force field and system-construction protocol differ from those used in the original AMBER/NAMD studies, the resulting free-energy surface and local barrier heights more closely resemble those reported in the recent NAMD tutorial than the original GaMD benchmark profiles.

### Folding of Chignolin

Starting from an extended conformation of chignolin, GaMD simulations implemented in GROMACS 2025.4 were able to capture complete folding of the protein into its native structure within 300 ns. The RMSD between the simulation-folded chignolin and the NMR experimental native structure (PDB: 1UAO)^12^ reached a minimum of 0.07 nm (Figure 2A). The boost potential applied in the GaMD simulations followed a Gaussian distribution, with an anharmonicity of 2.4 × 10^-3^ (Figure 2B). In three independent 300 ns GaMD simulations, chignolin reached the folded state (RMSD < 0.02 nm) in all three runs, and two of the simulations exhibited multiple folding-unfolding events. In sim 3, after remaining in the intermediate state (“I”) for 50 ns, the system stayed in the folded state thereafter (Figure 2C). During folding, the radius of gyration (*R_g_*) of chignolin decreased to 0.5 nm (Figure 2D). The average folding time of chignolin obtained from the GaMD simulations was approximately 100 ns, which is significantly shorter than the 600 ns folding time reported in previous long-timescale cMD simulation.^28^

**Figure 2.**
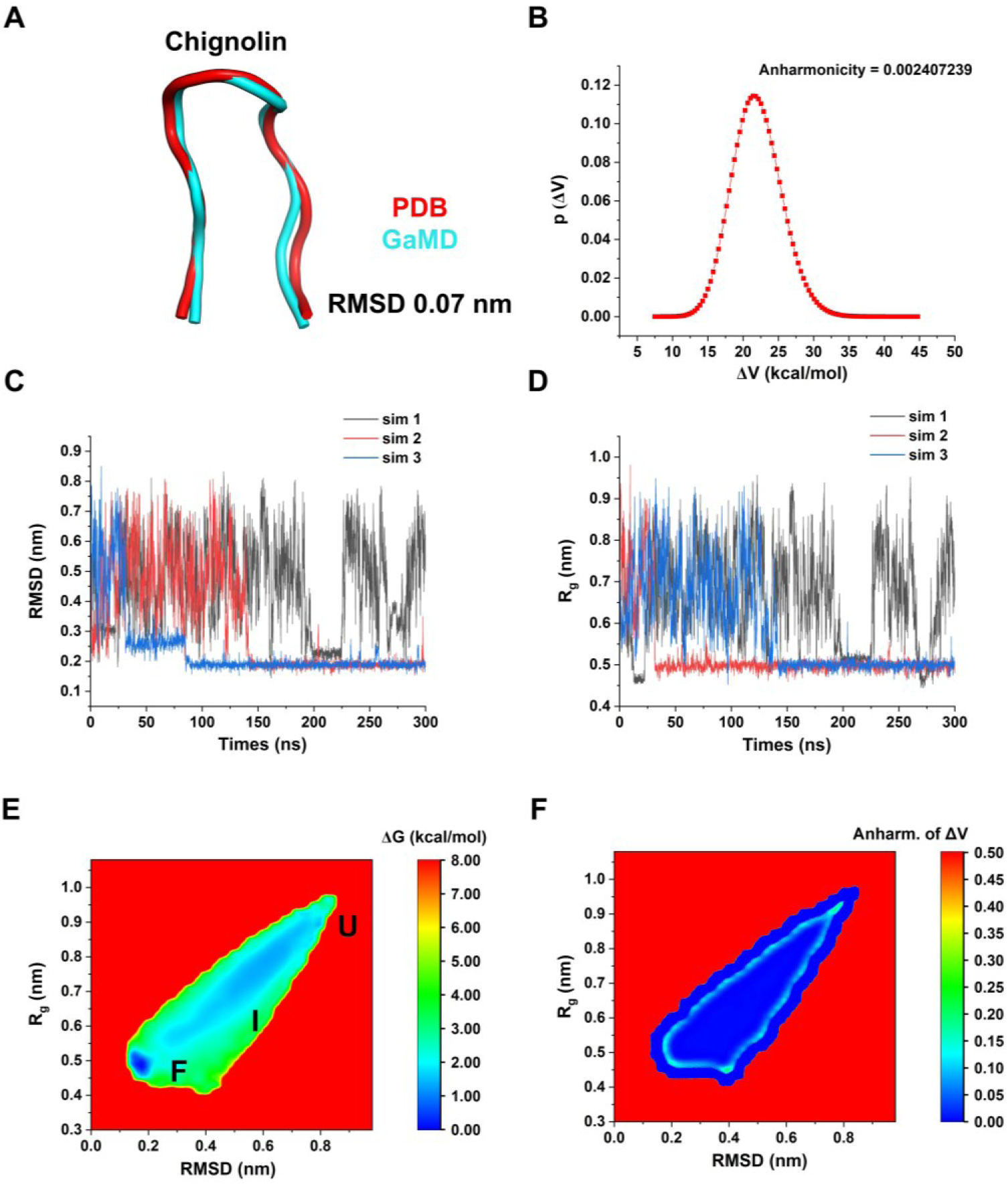
GaMD folding results of the chignolin system. (A) Superposition of the folded conformation sampled by GaMD and the reference PDB conformation. (B) Probability distribution of the boost potential (Δ*V*) during the 300 ns GaMD production stage, with an anharmonicity of 2.4 × 10^-3^. (C, D) Time evolution of the RMSD (C) and radius of gyration (*R_g_*, D) of chignolin in three independent 300 ns GaMD trajectories. (E) Two-dimensional reweighted PMF constructed based on RMSD and *R_g_* in which three major conformational basins can be resolved: folded (“F”), intermediate (“I”), and unfolded (“U”). (F) Local anharmonicity distribution of the boost potential on the (RMSD, *R_g_*) plane.

Based on the Gaussian distribution of the boost potential, cumulant expansion to the second order was applied to reweight the three combined 300 ns GaMD trajectories of chignolin. A two-dimensional PMF profile was then calculated using the protein RMSD relative to the native PDB structure and the radius of gyration (RMSD, *R_g_*), as shown in Figure 2E. The reweighted PMF allowed us to identify three distinct low-energy conformational states: the folded state (“F”), the unfolded state (“U”), and the intermediate state (“I”). Among them, the folded state corresponds to the global energy minimum located at (0.15 nm, 0.5 nm), while the unfolded state is located in a local energy well centered at (0.80 nm, 0.95 nm) and is 3.75 kcal/mol higher in energy than the folded state. The intermediate state spans a relatively broad region, with a large low-energy basin distributed over RMSD = 0.3–0.7 nm, and *R_g_* = 0.6–0.9 nm.

Figure 2F shows the distribution anharmonicity of Δ*V* for frames found in each bin of the two-dimensional PMF in Figure 2E. The anharmonicity remained below 0.1 throughout the conformational space sampled in the simulations. This indicates that the boost potential indeed follows a Gaussian distribution, thereby justifying the use of second-order cumulant expansion for reweighting. In summary, GROMACS-GaMD can effectively enhance sampling of the protein folding process and enable free-energy calculations, as demonstrated for the chignolin system.

### Folding of TC5b

Starting from an extended conformation, three independent 1 μs GaMD simulations successfully captured the folding of TC5b into native-like structures. The best-folded conformation closely overlapped with the reference PDB structure, with a minimum RMSD of 0.03 nm (Figure 3A). The boost potential showed a near-Gaussian distribution with an anharmonicity of 1.49 × 10⁻² (Figure 3B). All three trajectories reached the low-RMSD region, although folding occurred at different times (Figure 3C). The corresponding decrease in the radius of gyration to approximately 0.70 nm indicated the formation of compact conformations (Figure 3D). The experimental folding time of TC5b is approximately 3.6 μs at room temperature,^29^ while conventional MD studies have estimated a mean folding time of approximately 5.5 ± 3.5 μs.^30^ Therefore, the folding events observed within 1 μs in the present GaMD simulations indicate substantially enhanced conformational sampling.

**Figure 3.**
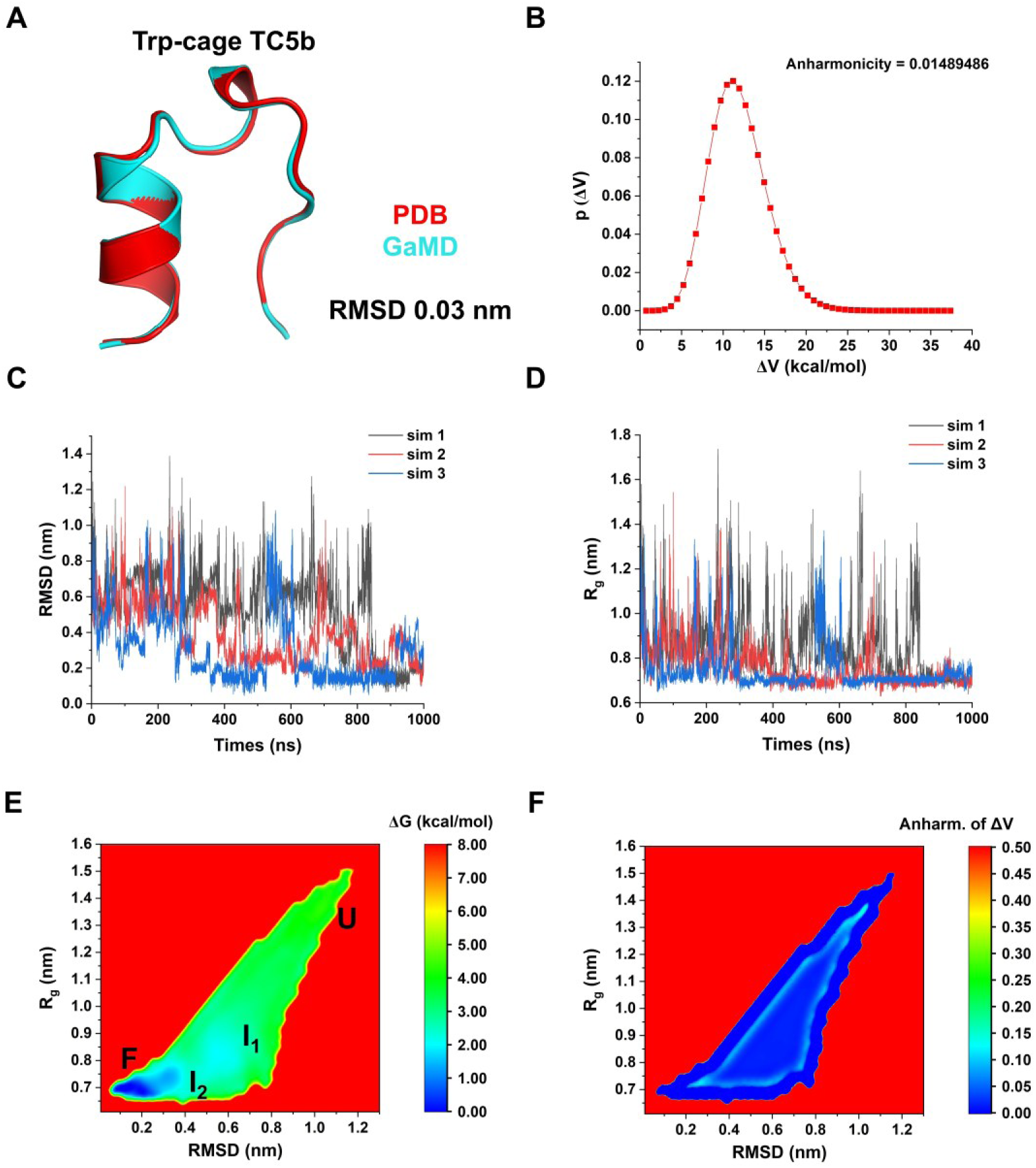
GaMD folding of TC5b. (A) Superposition of the best-folded conformation sampled by GaMD and the reference PDB structure. (B) Probability distribution of the GaMD boost potential (Δ*V*), showing an anharmonicity of 1.49 × 10^-2^. (C, D) Time evolution of the RMSD (C) and *R*_g_ (D) in three independent 1 μs GaMD simulations. (E) Two-dimensional reweighted PMF as a function of RMSD and *R*_g_, showing the folded (“F”), intermediate (“I₁” and “I₂”), and unfolded (“U”) conformational regions. (F) Local anharmonicity of the boost potential in the (RMSD, *R*_g_) space.

The combined trajectories were reweighted using second-order cumulant expansion to construct a two-dimensional PMF as a function of RMSD and *R*_g_ (Figure 3E). Four major conformational regions were identified, including the folded state (“F”), two intermediate states (“I₁” and “I₂”), and the unfolded state (“U”). The folded state corresponded to the global free-energy minimum at approximately RMSD = 0.1 nm and *R*_g_ = 0.7 nm. The intermediate I₂ was centered at approximately RMSD = 0.3 nm and *R*_g_ = 0.7 nm, whereas the more expanded intermediate I₁ occupied a broader region around RMSD = 0.5–0.7 nm and *R*_g_ = 0.8–0.9 nm.

Local anharmonicity remained below approximately 0.1 throughout most of the sampled conformational space (Figure 3F), supporting the reliability of the second-order cumulant expansion for free-energy reweighting.

### Binding of Benzene to T4 Lysozyme

In two out of five independent 500 ns simulations, the processes of benzene binding from the aqueous solvent into the ligand-binding site and dissociating from the binding site were sampled. After aligning the C-alpha atoms of T4 lysozyme, the minimum RMSD of the final benzene molecule relative to the bound conformation in the 181L^14^ X-ray crystal structure reached 0.06 nm (Figure 4A). The boost potentials applied during the two 500 ns GaMD simulations in which binding and unbinding events were observed followed a Gaussian distribution, with γ = 1.04 × 10⁻² (Figure 4B).

**Figure 4.**
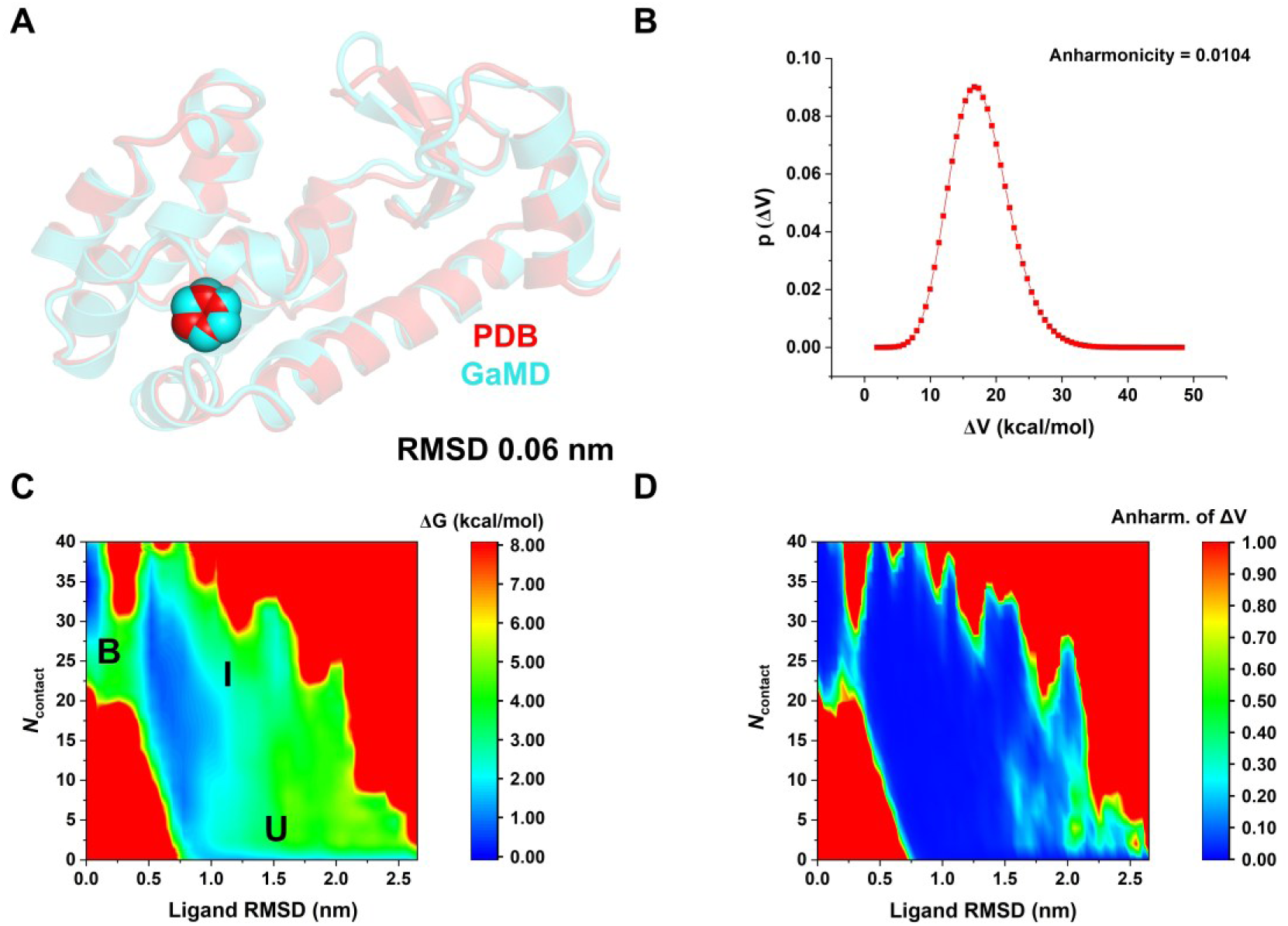
Binding of benzene to T4 lysozyme simulated by GaMD. (A) Comparison of the simulation-derived complex structure capturing benzene binding (cyan) with the 181L PDB structure (red), with a minimum ligand RMSD of 0.06 nm. (B) Distribution of the boost potential Δ*V* from two GaMD trajectories. (C) Two-dimensional (ligand RMSD, *N*_contact_) potential of mean force (PMF) calculated by reweighting the two GaMD simulation trajectories. (D) Distribution anharmonicity of Δ*V* for frames in each bin of the free energy profile.

Using the RMSD of benzene relative to the bound conformation and the number of protein heavy atoms within 0.5 nm of benzene (*N*_contact_), a two-dimensional PMF profile was calculated by reweighting the two 500 ns GaMD trajectories (Figure 4C). The reweighted PMF enabled us to identify three distinct low-energy states: the unbound state (“U”), the intermediate state (“I”), and the bound state (“B”). The bound state corresponds to the global energy minimum located at approximately (0 nm, 34); the unbound state lies in a local energy well centered at approximately (1.5 nm, 0); and the intermediate state features a relatively broad low-energy region centered at approximately (0.75 nm, 20). During the gradual entry from the aqueous solvent into the binding site, with *N*_contact_ progressively decreasing, the free energy gradually decreases, with an energy barrier of approximately 3.5 kcal/mol before entering the binding site (Figure 4C).

Figure 4D shows the anharmonicity γ of the Δ*V* distribution for the frames in each bin of the two-dimensional PMF. This value is relatively large in high-energy regions where sampling is limited. In contrast, in the energy-well regions, γ is less than 0.1, indicating that Δ*V* has been sufficiently sampled and is suitable for reweighting using a second-order cumulant expansion. Overall, Gromacs-GaMD can effectively enhance the sampling of ligand binding and unbinding processes within an acceptable simulation time and enable accurate free-energy calculations.

## DISCUSSION

This study implemented Gaussian accelerated molecular dynamics (GaMD) in GROMACS 2025.4 and validated its correctness, applicability, and enhanced-sampling capability using four representative systems. Unlike enhanced-sampling methods that rely on predefined collective variables, GaMD reduces effective energy barriers by applying a smooth harmonic boost potential to the potential energy surface, making it particularly suitable for complex biomolecular processes in which reaction coordinates are difficult to define a priori. Although GaMD has been supported to varying extents in platforms such as AMBER, NAMD, and OpenMM, a native implementation in GROMACS has long been lacking. This work fills this methodological and software-ecosystem gap, enabling GROMACS users to perform collective-variable-free GaMD enhanced-sampling simulations while retaining the advantages of high-performance parallel computing and GPU acceleration.

From an implementation perspective, integrating GaMD into GROMACS is not merely a straightforward port of an existing algorithm. The correct operation of GaMD requires real-time updating of potential-energy statistics, staged determination of boost parameters, consistency between the bias potential and biasing forces, reproducibility across restarts, and complete output for subsequent reweighting analysis. In this work, GaMD was embedded as an engine-level feature into the main MD workflow of GROMACS, with three modes implemented: total boost, dihedral boost, and dual boost. In particular, in the dual-boost mode, the program can simultaneously apply boost potentials to the total potential energy and the dihedral energy, thereby accelerating both global conformational fluctuations and local conformational transitions. By retaining GaMD parameters, energy statistics, boost potentials, and weight information required for reweighting, this implementation provides the necessary data foundation for subsequent free-energy reconstruction and trajectory analysis.

The results for the alanine dipeptide system show that this implementation can correctly reproduce the major features of the conformational free-energy landscape of a small molecule. After reweighting using the second-order cumulant expansion, the one-and two-dimensional PMFs obtained from 100 ns GaMD simulations were generally consistent with the 1000 ns conventional MD reference results, recovering the main low-energy conformational basins and the energy barriers associated with backbone dihedral rotations. This indicates that the calculation of the boost potential, force scaling, and reweighting output in the current implementation are internally consistent. At the same time, the two-dimensional free-energy landscape obtained in this study is not identical to earlier AMBER-GaMD or NAMD-GaMD benchmark results, mainly due to differences in force fields and system construction protocols. Earlier studies mostly used AMBER ff99-related parameters, whereas this study used the CHARMM36m force field and systems built with CHARMM-GUI.

In the chignolin folding system, GROMACS-GaMD sampled multiple transitions from unfolded to folded states within a 300 ns timescale and obtained folded conformations highly consistent with the experimental NMR structure. Compared with the folding time of approximately 600 ns reported in previous long conventional MD simulations, GaMD substantially shortened the simulation time required to reach the folded state in this study, demonstrating its ability to effectively lower free-energy barriers during folding and promote exploration of conformational space. The two-dimensional PMF constructed from RMSD and radius of gyration clearly distinguished the folded, intermediate, and unfolded states, indicating that this implementation not only accelerates conformational transitions but also provides a reasonable thermodynamic picture after reweighting.

The TC5b benchmark posed a more demanding test of the aforementioned capability because it is a larger 20-residue folding system with a more complex conformational energy landscape. Starting from extended conformations, all three independent 1 μs GaMD trajectories reached the low-RMSD folded region. The best-sampled structure exhibited a minimum RMSD of 0.03 nm relative to the experimental structure, indicating that the present implementation can recover a near-native folded structure even for a system substantially larger than chignolin. The observation of folding events within 1 μs is also consistent with the expected enhancement of sampling by GaMD: the experimental folding time of TC5b is approximately 3.6 μs at room temperature, whereas previous conventional MD simulations estimated a mean folding time of approximately 5.5 ± 3.5 μs. Consistent folding behavior was observed for both the chignolin and TC5b systems. All three independent trajectories for each system reached the folded state, while some trajectories additionally exhibited folding–unfolding round-trip events, further supporting the strong enhanced-sampling capability of GaMD for protein conformational transitions.

The benzene–T4 lysozyme system further demonstrates the applicability of this implementation to protein–ligand binding processes. Among five independent 500 ns GaMD trajectories, two sampled both the entry of benzene into the binding pocket and dissociation from the binding site, yielding bound conformations highly consistent with the crystal structure. The two-dimensional PMF constructed from ligand RMSD and the number of protein–ligand contacts distinguished the unbound, intermediate, and bound states, and revealed a distinct intermediate region and a finite binding barrier as benzene moves from the aqueous phase into the hydrophobic binding cavity. These results indicate that GROMACS-GaMD can enhance the sampling of ligand binding/unbinding events within an accessible simulation timescale and provide an effective tool for analyzing binding pathways and binding free-energy landscapes.

It is worth noting that the reliability of GaMD free-energy reweighting strongly depends on whether the boost-potential distribution is approximately Gaussian. In all four systems studied here, the overall boost-potential distributions were approximately unimodal and Gaussian, with low overall anharmonicity, supporting the use of second-order cumulant expansion for reweighting. In the alanine dipeptide, chignolin and TC5b systems, the local anharmonicity within the major low-energy basins remained low, indicating that the free-energy estimates in these regions are relatively reliable. In the benzene–T4 lysozyme system, the major energy basins, such as the bound and unbound states, also exhibited low anharmonicity. For these systems, higher anharmonicity was mainly observed in sparsely sampled high-energy regions. This result highlights an important principle in GaMD analysis: reweighted free energies in low-energy, well-sampled regions are generally more reliable, whereas interpretations of free energies in high-energy or low-probability regions should be made with caution.

Although this implementation demonstrates good accuracy and applicability, the current version still has several limitations. This work mainly implemented and validated the three standard GaMD modes—total boost, dihedral boost, and dual boost—and does not yet include selective or specialized GaMD variants designed for specific biophysical processes, such as LiGaMD, Pep-GaMD, and PPI-GaMD. Therefore, there remains room for further optimization when simulating specific systems.

In addition, the free-energy analyses in this study are still affected by the finite sampling length and the limited number of trajectories. For low-dimensional conformational systems such as alanine dipeptide, 100 ns GaMD was sufficient to recover the major free-energy features. For more complex processes such as protein folding and ligand binding, although GaMD significantly increased the probability of observing transition events, some intermediate states and high-energy regions may still be undersampled. In particular, in the benzene–T4 lysozyme system, binding/unbinding events were observed in only two trajectories. Therefore, the PMF of this system can be used to describe the main binding pathway and stable states, but further quantitative estimation of binding kinetics, dissociation rates, or standard binding free energies would require more independent trajectories, longer simulations, or integration with other kinetic analysis methods.

Future work may further extend the current implementation. Additional selective GaMD variants could be incorporated into GROMACS, such as ligand GaMD, peptide GaMD, and protein–protein interaction GaMD, to improve sampling efficiency for specific problems including ligand binding, peptide binding, and protein–protein recognition.

Overall, this study demonstrates the feasibility and effectiveness of a native GaMD implementation in GROMACS. Validation using four benchmark systems—alanine dipeptide, chignolin folding, TC5b, and benzene binding to T4 lysozyme—shows that the current implementation can stably generate approximately Gaussian boost-potential distributions, significantly enhance the sampling of conformational transitions and binding/unbinding events, and recover reasonable free-energy landscapes through second-order cumulant expansion. This work expands the enhanced-sampling capabilities of GROMACS, provides a new native tool for studying complex biomolecular conformational changes, protein folding, and ligand recognition processes, and lays the foundation for the future development of more efficient and more specialized GaMD methods.

## ASSOCIATED CONTENT

### Data and Software Availability

The source code and user tutorial for GROMACS-GaMD are available at https://github.com/math-diff/gromacs-gamd.

## AUTHOR INFORMATION

### Author Contributions

YY designed the experiments, performed the experiments, analyzed the data, and wrote the original manuscript.

### Notes

The authors declare no competing financial interest.

## ACKNOWLEDEMENTS

The author thanks Yinglong Miao for his helpful comments and suggestions on an earlier version of the manuscript.

